# Multiple forward scattering reduces the measured scattering coefficient of blood in visible-light optical coherence tomography

**DOI:** 10.1101/2022.03.20.485063

**Authors:** Raymond Fang, Ian Rubinoff, Hao F. Zhang

## Abstract

Optical properties of blood encode oxygen-dependent information. Noninvasive optical detection of these properties is increasingly desirable to extract biomarkers for tissue health. Recently, visible-light optical coherence tomography (vis-OCT) demonstrated retinal oxygen saturation (sO_2_) measurements using the depth-resolved spectrum of blood. Such measurements rely on differences between the absorption and scattering coefficients of oxygenated and deoxygenated blood. However, there is still broad disagreement, both theoretically and experimentally, on how vis-OCT measures blood’s scattering coefficient. Incorrect assumptions of blood’s optical properties can add additional uncertainties or biases into vis-OCT’s sO_2_ model. Using Monte Carlo simulation of a retinal vessel, we determined that vis-OCT almost exclusively detects multiple-scattered photons in blood. Meanwhile, photons mostly forward scatter in blood within the visible spectral range, allowing photons to maintain ballistic paths and penetrate deeply, leading to a reduction in the measured scattering coefficient. We defined a scattering scaling factor (SSF) to account for such a reduction and found that SSF varied with measurement conditions, such as numerical aperture, depth resolution, and depth selection. We further experimentally validated SSF in *ex vivo* blood phantoms pre-set sO_2_ levels and in the human retina, both of which agreed well with our simulation.

## Introduction

Optical coherence tomography (OCT) enabled noninvasive three-dimensional (3D) retinal imaging at micrometer-scale volumetric resolutions [1, 2]. Since its first report 30 years ago, OCT has become the clinical gold standard for diagnosing and monitoring nearly all major ocular diseases [3, 4].

OCT’s sensitivity to optical scattering and absorption provides imaging contrast between tissues and may be used to probe tissue health [5–8]. These capabilities extend to blood, where absorption and scattering are wavelength-dependent and oxygen-dependent [9–12]. Applying short-time Fourier transforms (STFTs), OCT can measure wavelength-dependent attenuations of blood with micrometer-scale depth resolution, enabling measurement of oxygen saturation (sO_2_) in discrete blood vessels [13–17]. Studies suggested that alterations in retinal sO_2_ can be a sensitive biomarker for blindness-causing diseases, including glaucoma and diabetic retinopathy [18, 19]. Hence, accurate and noninvasive retinal sO_2_ measurement can improve the clinical management of these diseases.

Blood’s optical absorption properties are 2-3 orders of magnitude higher in the visible spectral range than the near-infrared (NIR) spectral range [12], enabling recently developed visible-light OCT (vis-OCT) [14] to overcome the fundamental optical contrast limit in NIR OCTs [16, 20]. In 2013, Yi et al. showed that vis-OCT is sensitive to retinal sO_2_ in rodents. Later on, vis-OCT retinal oximetry was demonstrated in rodents [21–24] and humans [25, 26].

Vis-OCT relies on the reported optical properties of whole blood to estimate sO_2_. In the visible spectral range, Mie theory predicts an average absorption coefficient (*μ_a_*) near 150 cm^-1^ and an average scattering coefficient (*μ_s_*) near 3000 cm^-1^ [12, 27]. However, a wide range of experimentally measured values have been reported, suggesting uncertainty in the measured optical properties of blood, and potentially reducing the reliability of vis-OCT oximetry [9–11, 27–30]. For example, some measured *μ_s_* values are ~ 1/3 of the Mie theory prediction [9, 29, 31]. Researchers attributed such a reduction in *μ_s_* to the blood’s ‘packing factor’, which describes correlated optical interactions between densely packed RBCs and their hematocrit-dependence [9, 29]. Our group previously used the packing factor (denoted as *W*) to scale *μ_s_* in the vis-OCT inverse fitting model for sO_2_ measurement [31]. Specifically, our group found that the model’s goodness of fit (*R*^2^) maximized when *W* was between 0.2 and 0.4. This *W* value was consistent with the definition of the packing factor, which scales *μ_s_* by ~ 1/3 at physiological hematocrit [9]. Several other vis-OCT retinal oximetry works also used this *W* range to scale *μ_s_* [21, 25, 31, 32].

However, reported vis-OCT oximetry methods accounted for blood’s *μ_s_* differently, among which significant discrepancies exist [7, 22, 23, 31, 33–36]. Since vis-OCT oximetry fits the measured spectrum to the literature-reported *μ_a_* and *μ_s_*, deviations between the measured and reported *μ_s_* can introduce sO_2_ measurement error. Therefore, accurate and consistent quantification of sO_2_ benefits from a systemic investigation on how vis-OCT measures *μ_s_*.

We systemically investigated measuring optical scattering properties of blood detected by vis-OCT. First, we performed Monte Carlo (MC) simulations of a retinal blood vessel and photon detection by vis-OCT. MC simulation is not susceptible to systemic biases present in practical OCT detection or image reconstruction [37–39] and, therefore, provides a fundamental understanding of light-tissue interaction. For each photon packet exiting the tissue, we monitored the number of scattering events and optical pathlength traveled in tissue to investigate the impact of multiple scattering on the vis-OCT signal. We reconstructed simulated vis-OCT A-lines to establish a direct relationship between multiple scattering and the measured *μ_s_*. Then we established the scattering scaling factor (SSF), a generalized scaling coefficient for *μ_s_*. Since multiple scattering influenced the measured *μ_s_*, we further investigated photon detection by different numerical apertures (NA’s), because NA acts as a geometric filter in detecting multiply scattered photons. Second, we compared our simulation results to experimental vis-OCT imaging of *ex vivo* blood phantoms. We found excellent agreement between the simulated and experimentally measured values across pre-set oxygenation levels. Finally, we validated our SSF analysis in human retina vis-OCT imaging and found a strong agreement with our simulated results. Validated by simulation, *ex vivo* blood phantom imaging, and *in vivo* human retinal imaging, we provide evidence that vis-OCT measured *μ_s_* is smaller than the reported packing factor but higher than the reduced scattering coefficient 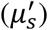 [37, 40]. This work sets the foundation for a unified theory of vis-OCT-measured optical properties of blood and more robust retinal oximetry.

### Detecting scattered light with vis-OCT

Most OCT’s use NIR (800 nm – 1300 nm) illumination, where biological tissues have lower optical scattering (*μ_s_* < 100 cm^-1^), lower optical absorbing (*μ_a_* < 1 cm^-1^), and moderately high scattering anisotropy (0.7 < *g* < 0.9) [41]. Such optical properties yield mean-free-paths (MFPs) of several hundred micrometers in tissue, meaning photons can travel deep into tissues before being multiply scattered. Since OCT’s axial resolution is dominated by light’s coherence length and not geometrical optics, OCT can use a low NA to image deeply penetrating photons across several hundred micrometers with axial resolutions < 10 *μ*m [42]. Another benefit of imaging weakly scattering tissues with a low NA is high sensitivity to single-scattered or ballistic photons [43]. In this work, we define ballistic photons as Class I photons [37], which satisfy

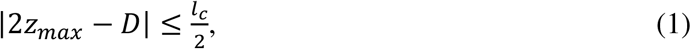

where *z_max_* [μm] is the maximum depth a photon travels in the tissue with respect to the OCT’s zero delay; *D* [μm] is the total optical path traveled by the photons with respect to the OCT’s zero delay; and *l_c_* [μm] is the coherence length. Essentially, a Class I photon travels straight, allowing only a slight deviation within *l_c_*. We define Class II photons as photons that do not travel in a straight line before being detected by the OCT, which satisfy

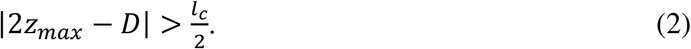

Finally, we define those photons undetectable by the OCT system as Class III photons.

Figs. 1A & 1B illustrate OCT imaging in a weakly scattering tissue (e.g. *μ_s_* < 100 cm^-1^ and 0.7 < *g* < 0.9) using a low NA (e.g. < 0.2). Fig. 1A shows a global illustration of tissue with optical properties within the NIR spectral range. The OCT sample arm focuses light on a spot in the tissue (illustrated by the green oval), which creates an A-line at that location. Due to the low NA, most incident photons (green arrow) are perpendicular to the tissue surface. Since the focus spot creates a conjugate point with the sample arm detector, it also acts as a geometric projection of the detector itself [44]. Only photons collected within the spot and the solid angle defined by the illumination NA will contribute to the A-line.

**Fig. 1.**
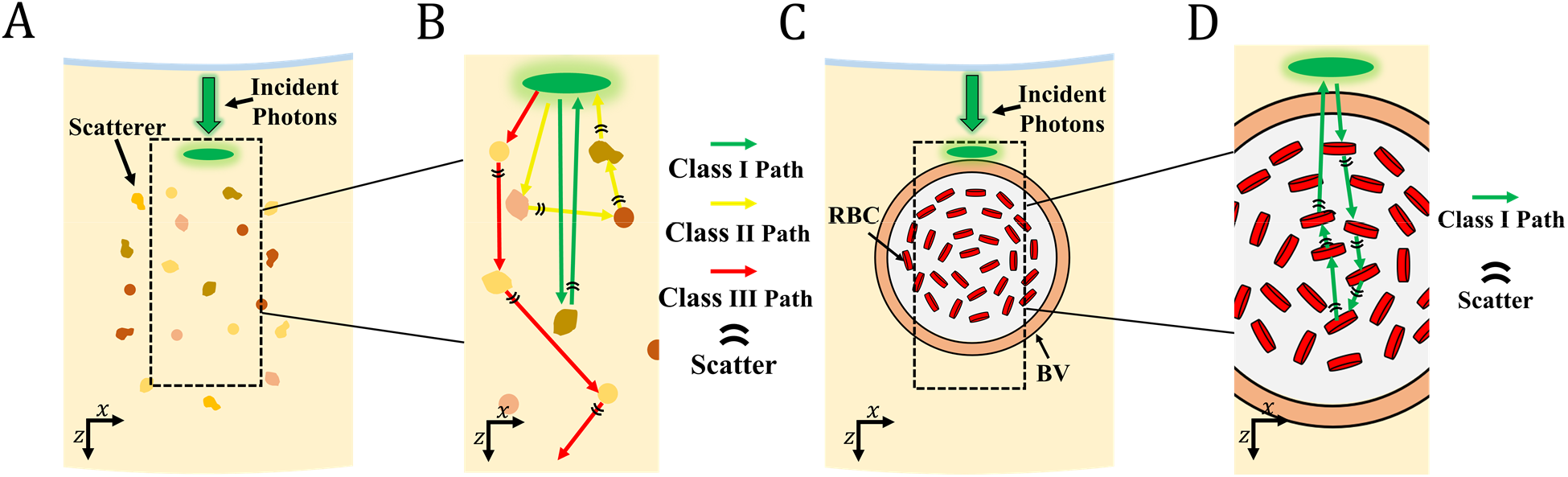
(A) Illustration of OCT illumination of a weakly scattering medium. Incident photons (green arrow) create a focal spot (green oval) and detection aperture in the tissue; (B) Magnified illustration of the highlighted region in the panel A, showing the paths of Class I (green arrow), Class II (yellow arrow), and Class III photons (red arrow); (C) Illustration of vis-OCT illumination of a retinal blood vessel (RBC) containing red blood cells (RBC); (D) Magnified illustration of the highlighted region in panel C, showing the paths of Class I photons (green arrow).

Fig. 1B is a magnified illustration of Fig. 1A in the black-dashed box. The green, yellow, and red arrows respectively illustrate representative paths of Class I, Class II, and Class III photons. The black bands represent scattering events. The Class I photon travels deeply into the tissue, scatters once, reverses direction, and returns along almost the same path in the *z*-direction (also referred to as backscattering). The Class II photons are scattered multiple times and travel significantly along the *x*-direction before returning to the detector. The Class III photons are scattered multiple times deeply in the tissue and do not return to the detector. Under the conditions of *μ_s_* < 100 cm^-1^ and 0.7 < *g* < 0.9, photons can travel tens or hundreds of *μ*m along the *x*-direction after each scattering event. Therefore, multiply scattered photons are increasingly likely to be Class III rather than Class II photons since they will travel too far from the detector before being absorbed by tissue or escaping the tissue. Meanwhile, most detected photons are likely to be Class I photons (green arrow in Fig. 1B) since they do not have the opportunity to travel outside the detection region.

When most photons are Class I and singly scattered, the OCT A-line at particular depth *z* can be modeled by the Beer-Lambert Law [45]

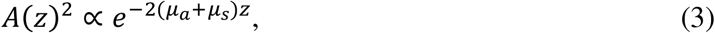

where *A*(*z*)^2^ [arbitrary unit] is the A-line intensity; *z* [mm] is the depth in the tissue; and the coefficient 2 in the exponential term indicates round trip in tissue. Eq. 3 has been thoroughly derived and experimentally validated for Class I photons [6].

Vis-OCT imaging of blood represents a special scenario that deviates from the assumptions described in Figs. 1A & 1B. The reported *μ_s_* for oxygenated and deoxygenated blood states are > 3000 cm^-1^ and the *W*-scaled *μ_s_* are near 1000 cm^-1^ [9]. Both these *μ_s_* values are over an order of magnitude greater than *μ_s_* values (< 100 cm^-1^) typical of most tissues within NIR spectral range. Furthermore, blood is more highly forward scattering (*g* ≥ 0.98) than most tissues (0.7 < *g* < 0.9). Therefore, photons are likely to travel only a few *μ*m or less along the *x*-direction after each scattering event. Assuming normal incidence of light on a vessel (Fig. 1C), photons can be multiply scattered and still satisfy the Class I condition. Fig. 1D illustrates such a path following the green arrow. We hypothesize that the path shown in Fig. 1D is a common, if not dominant, detection scenario in blood imaging using vis-OCT. A Class I photon that travels deeper than its single scatter assumption is equivalent to reducing its *μ_s_* in Eq. 3. Therefore, the Beer-Lambert Law may be rewritten as

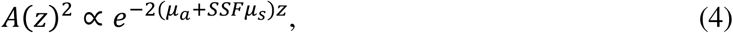

where SSF [dimensionless] is the scattering scaling factor, which is < 1 and scales *μ_s_* to account for the increased detection of photons deeper in tissue. The SSF is a generalized scaling factor and incorporates any other reductions of *μ_s_*, including W.

### Vis-OCT oximetry

Oxygenated and deoxygenated blood have distinct wavelength-dependent *μ_a_* and *μ_s_*, allowing estimation of sO_2_. An STFT [46] can reconstruct spectrally-dependent A-lines, which can be modeled by

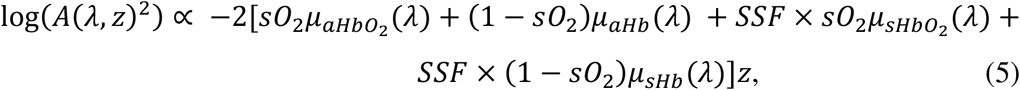

where *λ* [nm] is the central wavelength of the selected STFT sub-band; sO_2_ [dimensionless, between 0 and 1] is the oxygen saturation in blood; the subscripts HbO_2_ and Hb denote oxygenated or deoxygenated hemoglobin, respectively; and SSF [dimensionless] is the scattering scaling factor. We used 21 STFT sub-bands ranging from 528 nm to 588 nm equidistant in wavenumber, with an average full-width-at-half-max (FWHM) bandwidth of 11 nm.

## Methods

### MC simulation parameters

We simulated vis-OCT detection and reconstruction in a retinal blood vessel using MC simulation [8, 27, 37, 40, 47–51]. Fig. 2 shows the multi-layered 3D model of a blood vessel embedded in the retina. We modeled the blood vessel using an infinitely long cylinder located 55 *μ*m below the vitreous-retina interface. The vessel has three concentric layers: blood, cell-free zone (CFZ) [35, 36], and the vessel wall. The CFZ is a thin layer consisting primarily of plasma between the blood and the vessel wall. It arises from the difference in viscosity between RBCs and plasma and is described by the Fahraeus Lindqvist (FL) effect [52]. The CFZ has been previously observed in OCT images [53] and is noticeable in our vis-OCT data. Table I summarizes the optical and geometrical parameters. We extracted the properties of the retina [27, 49], vessel wall [27], and CFZ [54] from the literature. We used the theoretical optical properties of blood previously derived by Faber et al. [12, 27]. To increase the speed of our simulation, we used the average *μ_a_* and *μ_s_* between 520 nm and 600 nm (center wavelength 560 nm). Although there is still uncertainty in the exact values of *μ_a_, μ_s_*, and *g* [9], our values are well within the reported range [9–11, 27–30]. To account for correlated optical interactions between densely packed RBCs, we scaled blood’s *μ_s_* using the packing factor 4 = (1 − *H*)^2^, where *H* is hematocrit [dimensionless]. Assuming a hematocrit of 45% [55], we have *W* = 0.3025

**Fig. 2.**
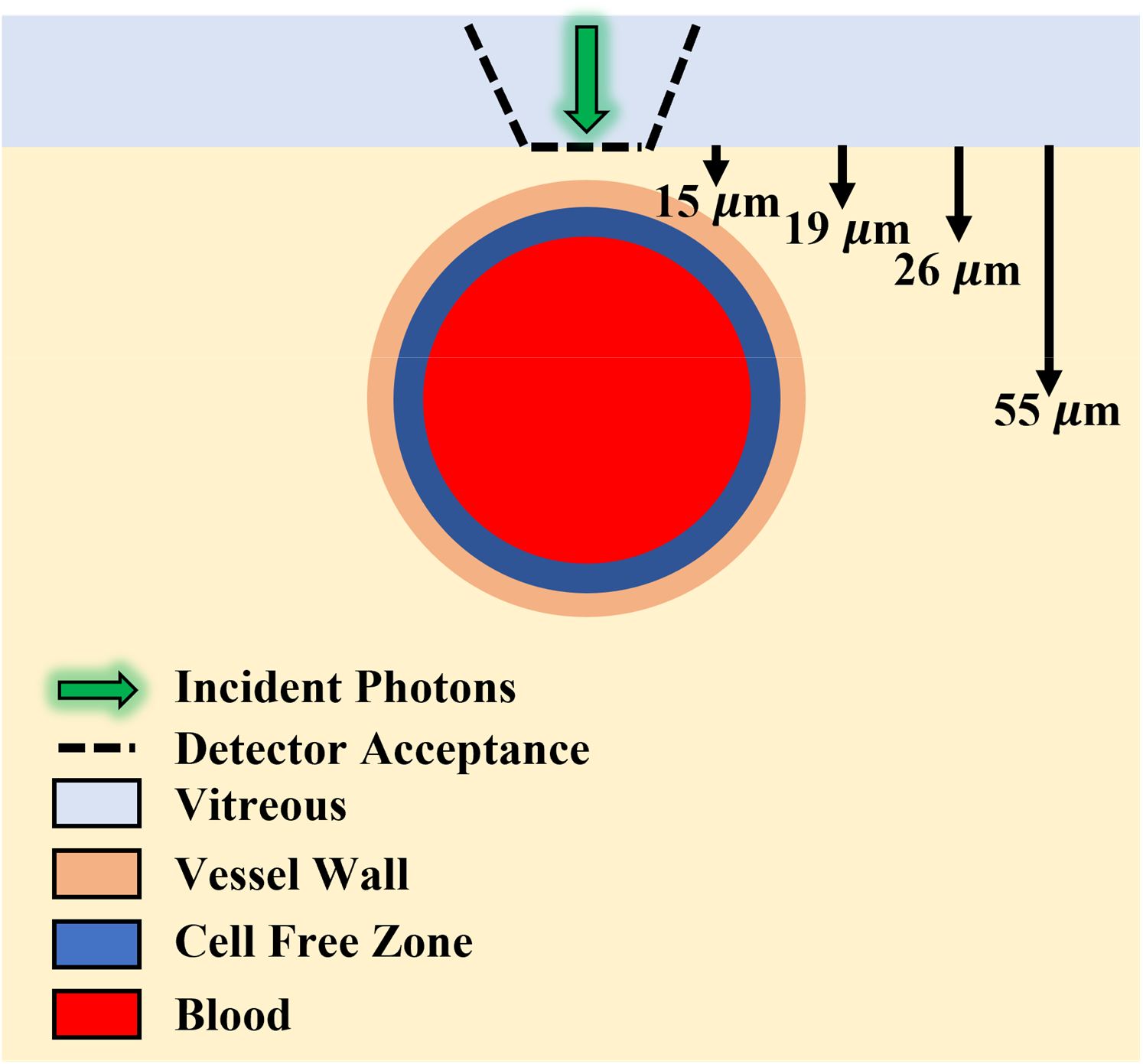
Illustration of tissue and imaging geometry in Monte Carlo simulation. Downward arrows (from left to right) highlight the interface depths of the vessel wall, cell free zone, and blood, respectively; fourth arrow highlights depth of the vessel center

### MC simulation algorithm

MC simulation of photon propagation in biological tissue has been widely reported [8, 27, 37, 40, 47–51, 56]. We followed the algorithm of simulation photon transport in multi-layered tissues (MCML) [40]. Briefly, we launch a photon packet towards the retina and blood vessel, as illustrated in Fig 2. The photon packet’s launching position is at the vitreous-retina interface (Fig. 2) at the lateral center of the vessel. The initial direction vector is along the *z*-axis. Upon tissue entry, we generated the stepsize (*s*) following the Poisson distribution

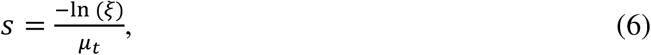

where *μ_t_* = *μ_a_* + *μ_s_* and *ξ* [dimensionless] is a random variable following an even distribution between 0 and 1. After traveling distance *s*, the photon packet interacts with tissue and deposits a fraction 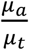 of its weight. At each interaction location, the photon packet is scattered by an angle *θ* relative to its current propagation direction determined by the Henyey-Greenstein phase function [56]

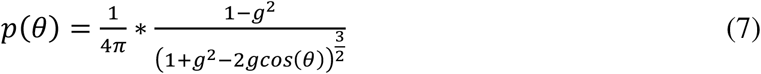

At the interface between two media, the photon packet either reflects or transmits with probabilities according to the Fresnel’s equations [57]. Upon entering a new tissue region, *μ_t_* is adjusted accordingly. The simulation continues until the photon packet exits the retina into the vitreous or the optical path distance traveled is greater than 2000 *μ*m, well beyond the depth of the blood vessel.

If the photon packet exits the retina into the vitreous, we recorded the remaining weight of the photon packet, the optical pathlength traveled within the tissue, the total number of scattering events in all tissue regions, the total number of scattering events in blood, the exiting position, and the exit angle. We simulated 10^10^ photon packets for each A-line. We implemented the simulation in MATLAB 2020 using parallel computing on a PC with a 3.4-GHz Intel Core i7-6800K CPU and 64-GB RAM. The simulation of an A-line took approximately 120 hours to complete.

### Photon detection

To simulate OCT detection, we geometrically filtered photons exiting the retina [51] (black-dashed lines in Fig. 2). We determined photon acceptance aperture and angle according to the NA of the light incident on the retina. We tested the NA value from 0.015 to 0.094, which follows the optical properties of normal human eyes [58]. We calculated the NA as

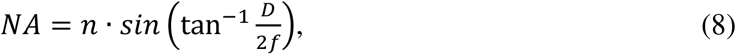

where *n* [dimensionless] = 1.35 is the refractive index; *D* [cm] is the 1/*e*^2^ diameter of the collimated beam incident on the cornea; *f* [cm] = 1.8 cm is the focal length of a normal eye [59]; and 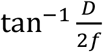 is the acceptance angle with respect to normal incidence. We used the NA to calculate the focal spot beam waist

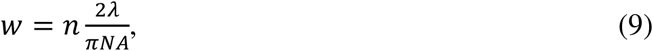

where *λ* = 560 nm is the central wavelength of the vis-OCT probing light. For simplicity, we detect photons using a uniform circle with a radius of *W*. In the results, we use an NA = 0.05, equivalent to a 2.9° acceptance angle and a 7.0 *μ*m detection aperture diameter unless otherwise specified.

### Vis-OCT A-line reconstruction

We reconstructed simulated vis-OCT A-lines using the recorded photon weights and optical path distances in the simulation. We only used photons detected under the acceptance conditions. Adopting the methods in Kirillin et al. [51], we reconstructed the OCT A-line as

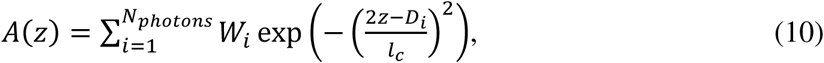

where *W_i_* [dimensionless] is the photon weight; *D_i_* [cm] is the optical path distance; and *l_c_* [*μ*m] is the axial resolution. We set the distance between adjacent *z* positions as 1 *μ*m and used an axial resolution of 9 *μ*m defined by the *l_c_* of the STFT sub-band window centered at 558 nm with an FWHM bandwidth of 11 nm [42]. We used this sub-band window size for spectroscopic A-line reconstruction in our experimental studies.

### Experimental measurements of *ex vivo* blood samples

We used the vis-OCT system operating from 510 to 610 nm described by Beckmann et al. [60] to image *ex vivo* blood samples in phantom vessels. The imaging objective in the sample arm had an NA of 0.05 [61], consistent with our simulation and human imaging. Briefly, we constructed a vessel phantom by pulling a glass capillary tube to have an inner diameter of 200 *μ*m and embedded and stabilized the tube in a plastic well. To reduce the influence of specular reflections, we added immersion oil to the well until the tube was covered. We prepared whole bovine blood (Quadfive, Ryegate, MT) of hematocrit 45% of oxygen levels ranging from 45% to > 99%. To oxygenate the blood, we added a constant stream of pure oxygen and stirred the blood with a magnetic stir bar. To deoxygenate the blood, we added sodium dithionite [62] to the solution and stirred. We repeated these processes until reaching the desired oxygen level. We monitored blood’s partial pressure of oxygen (pO_2_), partial pressure of carbon dioxide (pCO_2_), pH, and temperature using a blood-gas analyzer (Rapidlab 248, Siemens Healthcare Diagnostics, Malvern, PA) and estimated the corresponding sO_2_ [63]. Before loading the tube with blood, we flushed it with a phosphate-buffered saline and heparin solution to reduce clotting or sedimentation. Finally, we loaded the tube with blood and flowed it at 0.3 mm/s using a syringe pump (Fusion 100, Chemyx, Inc. Stafford, TX). We aligned the tube to be in focus and near the system zero-delay and illuminated it with 1.20 mW of power. Finally, we acquired data consisting of 512 A-lines × 256 B-scans with imaging range 1 mm × 1 mm at a 25 kHz A-line rate.

### Experimental measurement of human retinal vessels

For human imaging, we used the system described by Rubinoff et al. [64]. We used Eq. 8 to estimate an NA of 0.05 in the retina, similar to the simulation and *ex vivo* measurements. All human imaging procedures were approved by Northwestern Institutional Review Board (IRB) and adhered to the Tenets of Helsinki. We illuminated the retina with 250 *μ*W of power and acquired human retinal images consisting of 8192 A-lines × 16 B-scans repeated across a 3.8 mm field of view at a 25 kHz A-line rate.

## Results

### Contribution from multiple forward scattering in vis-OCT blood signal

We investigated how blood’s scattering properties influence the detection of photons in vis-OCT. Fig. 3A plots a detected photon packet path from our MC simulation. The concentric rings plot the outer boundaries of each vessel layer. The photon packet launched from the origin followed the path of the green line. The green asterisk (*) highlights a scattering event, and a circled asterisk highlights a transmission or reflection across tissue regions. The photon packet in Fig. 3A was scattered 11 times inside the blood vessel. Notably, the photon packet travels mainly along the *z*-axis, consistent with a high scattering anisotropy (g = 0.987). This allows the photon packet to backscatter to nearly the same *x*-position as it launched, similar to the illustration in Fig. 1D. Despite the multiple scattering events, we classified this photon packet to be Class I. Based on the optical properties shown in Table 1, the calculated MFP in blood was 8.5 *μ*m; however, the photon packet travels 60 *μ*m into the vessel (Fig. 3A).

**Fig. 3.**
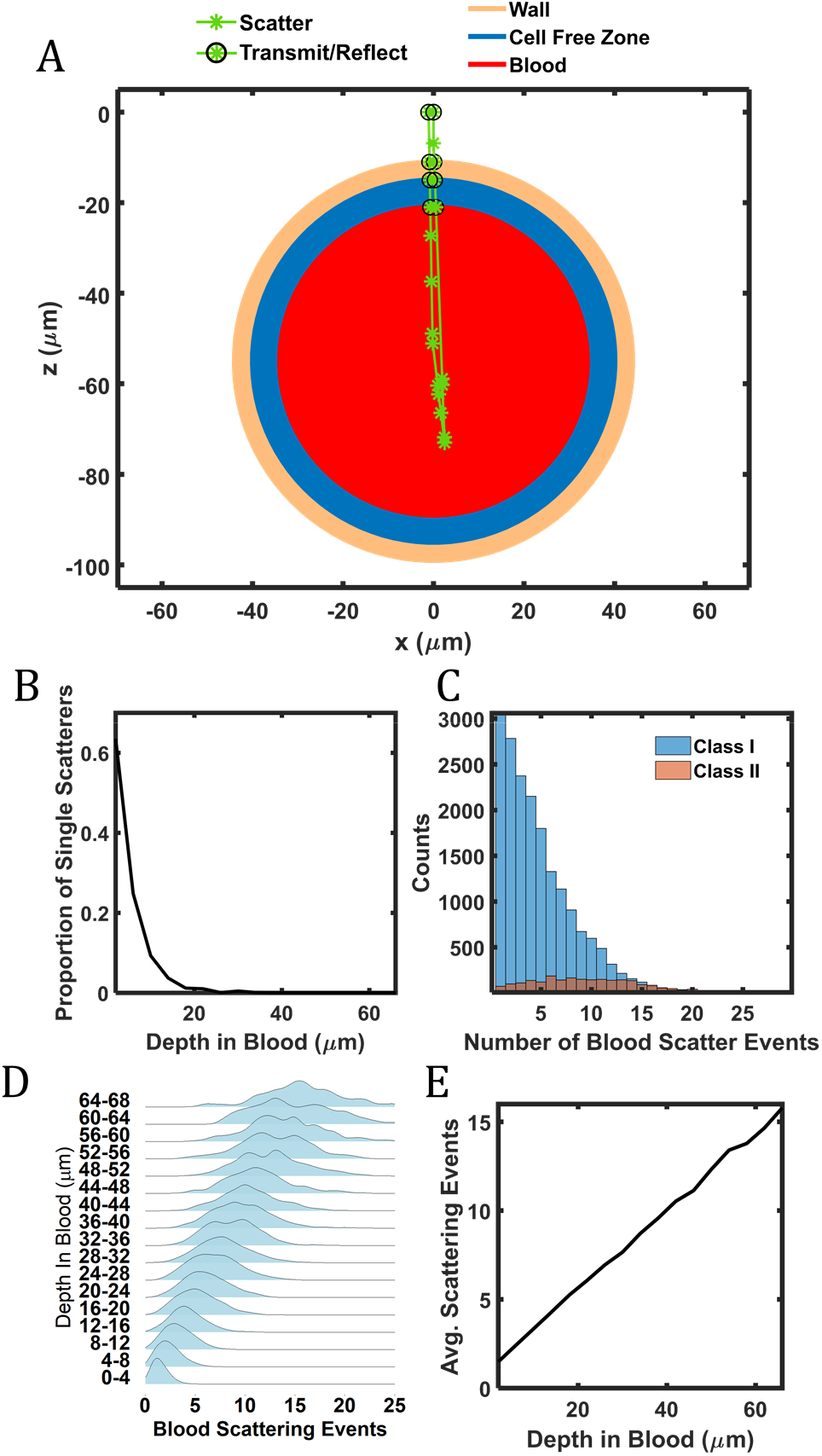
(A) Simulated photon packet path (green line) and scattering events (green stars) in the blood vessel; (B) Proportions of detected singly-scattered photon packets; (C) Histogram showing the number of scattering events of the detected Class I and Class II photons; (D) Distributions of the number of scattering events at different depths in blood; (D) The average number of scattering events at different depths in blood.

**Table 1.**
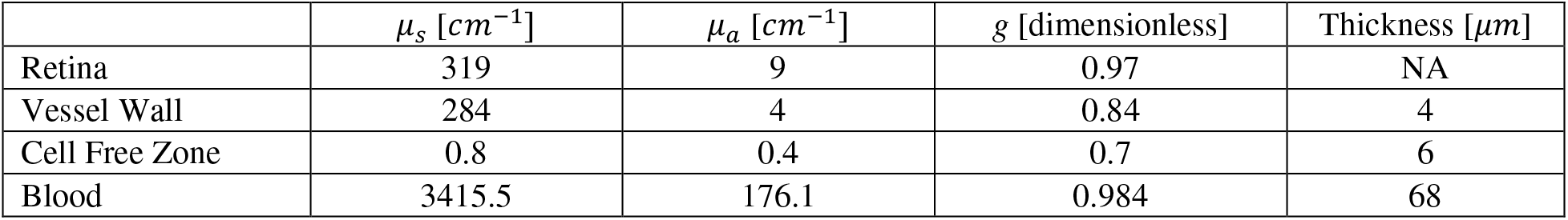
Optical properties of tissues used in Monte Carlo simulation.

To assess whether the multiple scattering observed in Fig. 3A was a frequent occurrence, we measured the proportion of all detected singly scattered photons packets that entered the blood region (Fig. 3B). Within the first 10 *μ*m in blood, about 50% of photon packets are singly scattered, which is consistent with the predicted MFP near 8.5 *μ*m. The proportion drops rapidly, where almost no detected photons are singly scattered after 17 *μ*m into the blood region (2 MFPs). Considering that measurements deeper than 8.5 *μ*m are necessary to provide sufficient attenuation contrast, it is reasonable to assume that retinal oximetry measurements are dominated by multiply scattered photons (excluding capillaries that may not generate multiple scattering due to low RBC counts).

Fig. 3C plots the detected Class I and Class II photons that entered the blood region. To improve the precision of the photon packet classification, we used *l_c_* = 1.4 *μ*m, the estimated vis-OCT full band resolution, rather than *l_c_* = 9 *μ*m, the STFT resolution. As shown in Fig. 3C, Class I photons dominate. Figs. 3B & 3C collectively demonstrate that most detected photons are both Class I and multiply scattered photons. This observation is further emphasized by Fig. 3D, which plots the fitted histograms of scattering events in blood from detected photons for different depths. The histograms are normalized with respect to their maximum values to show detail. The distributions show that, other than at the shallowest depths, photons are always multiple scattered. The average value of scattering events increases, and their distributions broaden as photon packets propagate deeper. Fig. 3E shows that the average value of scattering events is always > 1 and increases linearly with depth. Such a linear increase is consistent with the notion that a photon packet travels nearly along the same direction after each scattering event. The slope of this line corresponds to 1 blood scattering event for a photon step of 8.9 *μ*m, consistent with the calculated MFP.

It is commonly believed that Class I photons are singly scattered and Class II photons are multiply scattered in OCT tissue imaging. Fig. 3 suggests that photon propagation in the blood is similar to a scenario where singly scattered photons are detected from a medium with a significantly smaller *μ_s_*. This is why we introduce the scattering scaling factor SSF for *μ_s_* in the modified Beer-Lambert model in blood as defined in Eq. 4.

### Measuring the SSF value

Fig. 4A plots the simulated A-line on a natural logarithm scale (referred to as ‘log’). and illustrates the depth selection procedure for SSF measurement. From left to right, the first peak represents the anterior wall (AW) of the vessel, which is slightly convolved with the retinal tissue above it. Beneath the AW is a valley representing the cell-free zone (CFZ). The valley is smaller than expected from a completely ‘scattering-free’ region due to the limited 9-*μ*m axial resolution. Beneath the CFZ is the blood maximum (BM), representing the start of blood signal decay (BSD) in the A-line. The distinct peak at the BM is collectively contributed by the finite MFP in blood, the size of the CFZ, and the limited 9-*μ*m axial resolution. The depth location of BM in the A-line is about 10 *μ*m deeper than the physical start of the vessel lumen and about 20 *μ*m deeper than the AW peak. The blood signal decay (BSD) follows the BM and is consistent with log-scale decay described by the Beer-Lambert law [45]. When measuring the optical properties of blood *in vivo*, it is critical to start at least from the BM rather than assuming BSD occurs at the boundary of the AW and vessel lumen. Finally, the last peak represents the posterior wall (PW) of the vessel.

**Fig. 4.**
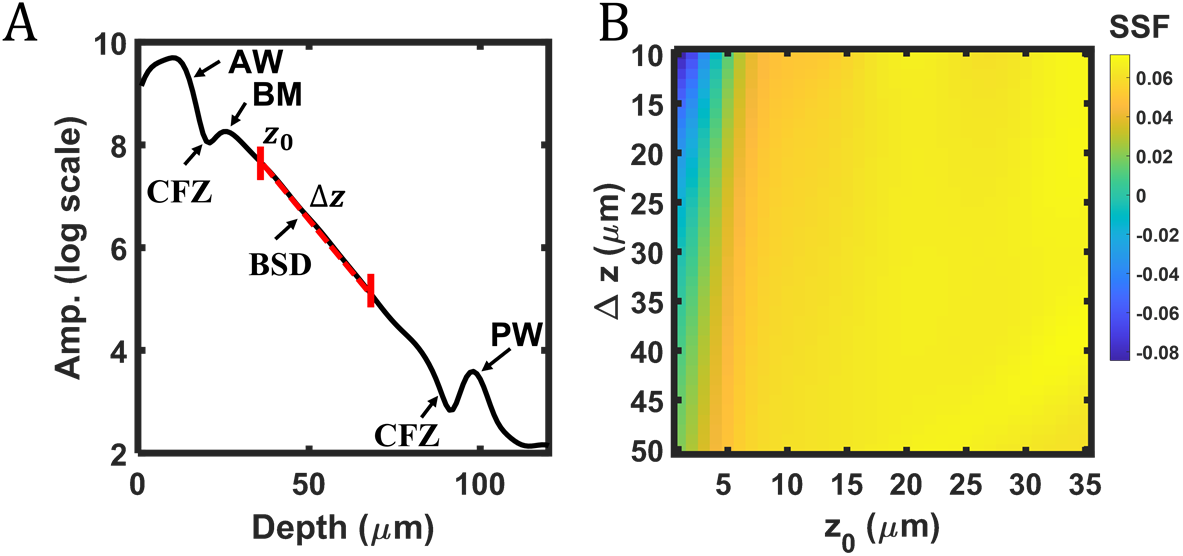
(A) A simulated A-line at the center of the vessel, showing the anterior wall (AW), cell-free zone (CFZ), blood maximum (BM), blood signal decay (BSD), and posterior wall (PW); The red-dashed line highlights measurement depth range starting at *z*_0_ and ranging Δ*z*; (B) Measured SSF values from A-line in the panel A as a function of different *z*_0_ and Δ*z*.

To measure the scattering coefficient, we modeled the BSD using a modified Eq. 5

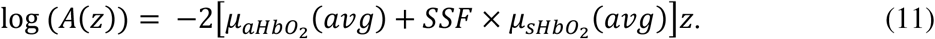

We include only the oxygenated coefficients since the simulated sO_2_ was 100%. The term ‘avg’ indicates that simulation used optical properties that were the average value between 520 nm and 600 nm. The double pass Beer-Lambert law does not need to be squared in the simulated A-line since it is not the product of interference between the sample and the reference electromagnetic fields. Although there are a handful of ways to extract *μ*_*sHbO*_2__(*avg*) from this equation, we elected to compute a depth-average of Eq. 11, which we empirically found robust against noise [64]. The starting measurement depth is *z*_0_ and the depth range is Δ*z*. The full region of measurement is highlighted by the red dashed line. We first normalized *A*(*z*) by its amplitude at *z*_0_, which shifted the coordinate system to *z*_0_ = 0. The average intensity becomes

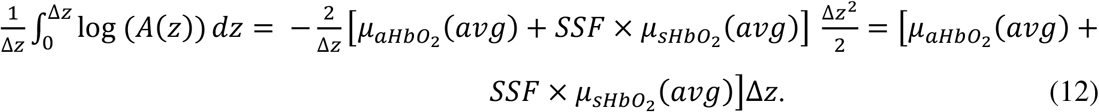

Dividing by Δ*z*, we are left only with the linear combination of *μ*_*aHbO*_2__(*avg*) and *SSF* × *μ*_*sHbO*_2__(*avg*). By subtracting *μ*_*aHbO*_2__(*avg*), whose value is from the literature, we are left with *SSF* × *μ*_*sHbO*_2__(*avg*), which can be compared with the literature *μ_s_* value used in the simulation. The SSF can be calculated as

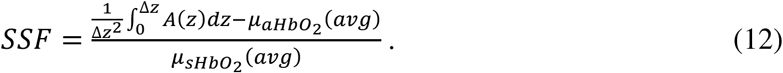

Fig. 4B shows the SSF under various combinations of *z*_0_ and Δ*z*, where *z*_0_ = 0 at the start of the vessel lumen. For *z*_0_ between 0 *μ*m and 5 *μ*m, SSF ≤ 0, which is caused by choosing a depth before BM and is physically impossible. At *z*_0_ = 10 *μ*m, the approximate location of BM, SSF = 0.05-0.06, with slight variation for different Δ*z*. For *z*_0_ between 11 *μ*m and 25 *μ*m, SSF = 0.060-0.068. For z0 > 25 *μ*m, SSF is slightly greater than 0.07, which may be biased by the CFZ near the PW. Therefore, we selected *z*_0_ = 17 *μ*m and Δ*z* = 33 *μ*m, a range where the SSF = 0.064. The measured SSF is approximately 5-fold smaller than the literature packing factor *W* = 0.3025. Here, *μ*_*sHbO*_2__ = 0.064 × *μ*_*sHbO*_2__(*avg*) = 222 cm^-1^, which is similar to the absorption coefficient *μ*_*aHbO*_2__(*avg*) = 174 cm^-1^. Considering *μ*_*sHbO*_2__(*λ*) has components that are also absorption dependent by the Kramers-Kronig relationship [9], we conclude that the spectrum of blood vis-OCT measures is overwhelmingly dominated by absorption instead of scattering.

### Influence of the number of scattering events on SSF

To understand how multiple scattering influences SSF, we set a graded threshold to the number of detected scattering events in blood. If a photon packet that entered the blood region is scattered more times than the set threshold, it was not included in the simulated A-line. Fig. 5A shows the simulated A-line for different scattering thresholds. The different shades of red plot different scattering threshold levels. The darkest shade has no threshold and collects photon packets from all scattering events (same as Fig. 4A). A-line amplitude decays slower with higher threshold levels, consistent with the notion that increased multiple scattering reduces SSF. Specifically, when the threshold level is 1, the A-line is reconstructed by singly-scattered Class I photon packets and decays within 20 *μ*m in blood. As sO_2_ calculation fits A-lines beyond 20 *μ*m in blood, multiply scattered photons are required for accurate sO_2_ measurement *in vivo*.

**Fig. 5.**
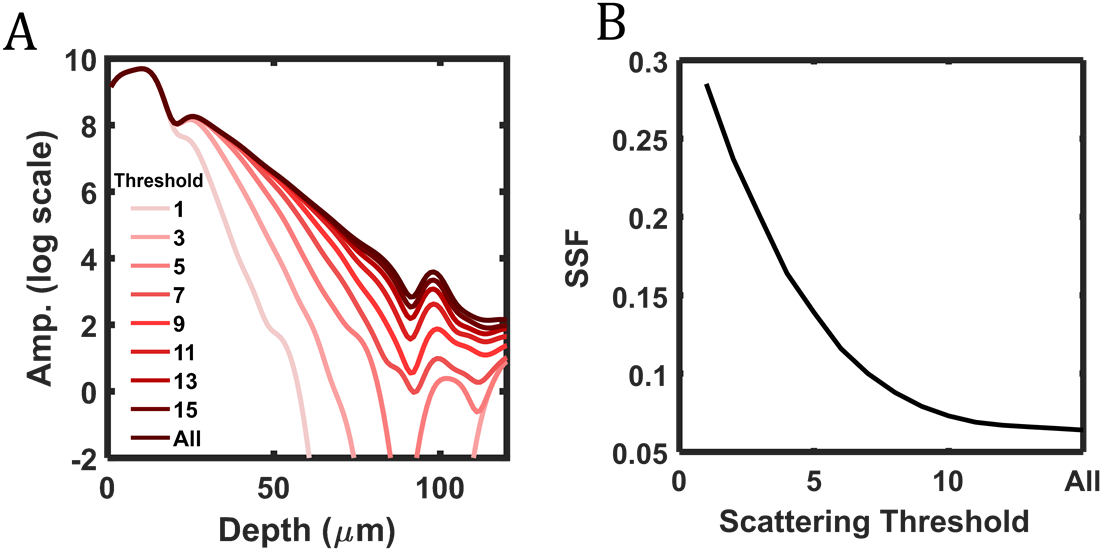
(A) Simulated A-line at the center of the vessel after thresholding the number of detected scattering events in blood. Lighter shades indicate a lower threshold of detected scattering events; (B) The SSF value as a function of detected scattering threshold.

Additionally, it becomes clear that multiple scattering enables deeper photon penetration necessary for visualizing the PW. The PW becomes weak when the threshold level is less than 7 and invisible when the threshold level is less than 3. This suggests that multiple forward scattering also facilitates the visualization of the PW, a landmark commonly used in vis-OCT oximetry to indirectly measure *μ*_s_ [31]. Fig. 5B shows the relationship between SSF and the scattering threshold level, which decays approximately exponentially with the scattering threshold, asymptotically approaching 0.064 after threshold > 15. Thresholding 7 scattering events gives SSF = 0.116 and thresholding 2 scattering events gives SSF = 0.285, approaching the set *W* = 0.3025. We measured SSF = 0.424 when thresholding 1 scattering event, although we did not include this data point due to insufficient signal-to-noise-ratio along the measured depths.

### Influence of numerical aperture on SSF

As described in photon packet detection in our MC simulation, the detection NA acts as a spatial filter for photon scattering events. The detection aperture (radius) and angle limit an existing photon packet’s position and propagation direction. Therefore, the detection criteria can potentially influence the measured SSF.

We examined the relationship between NA and SFF for *z*_0_ = 17 *μ*m and Δ*z* = 33 *μ*m based on the conclusion from the above section. The tested NA’s are 0.015 to 0.094, which are based on physically reasonable imaging NA’s in the human eye. In OCT, retinal imaging NA is almost always less than the maximum possible NA (e.g. 0.2 [58]), since researchers must consider limiting factors like reduced depth-of-focus, aberrations, and eye dilation [44]. Therefore, we varied the acceptance radius from 1.9 μm to 11.9 μm and varied the acceptance angle from 0.9° to 5.4° and found that SSF varied between 0.02 to 0.09 (Fig. 6A). We found that SSF reduces with increased acceptance radius, allowing more photon Class II photons (result here). We also found that SSF increased with increased acceptance angles. Photon packets traveling deeper in tissue are less likely to be filtered by the acceptance angle as most photon packets within the acceptance radius for deeply traveling photon packets are within the acceptance angle in deeper tissues.

**Fig. 6.**
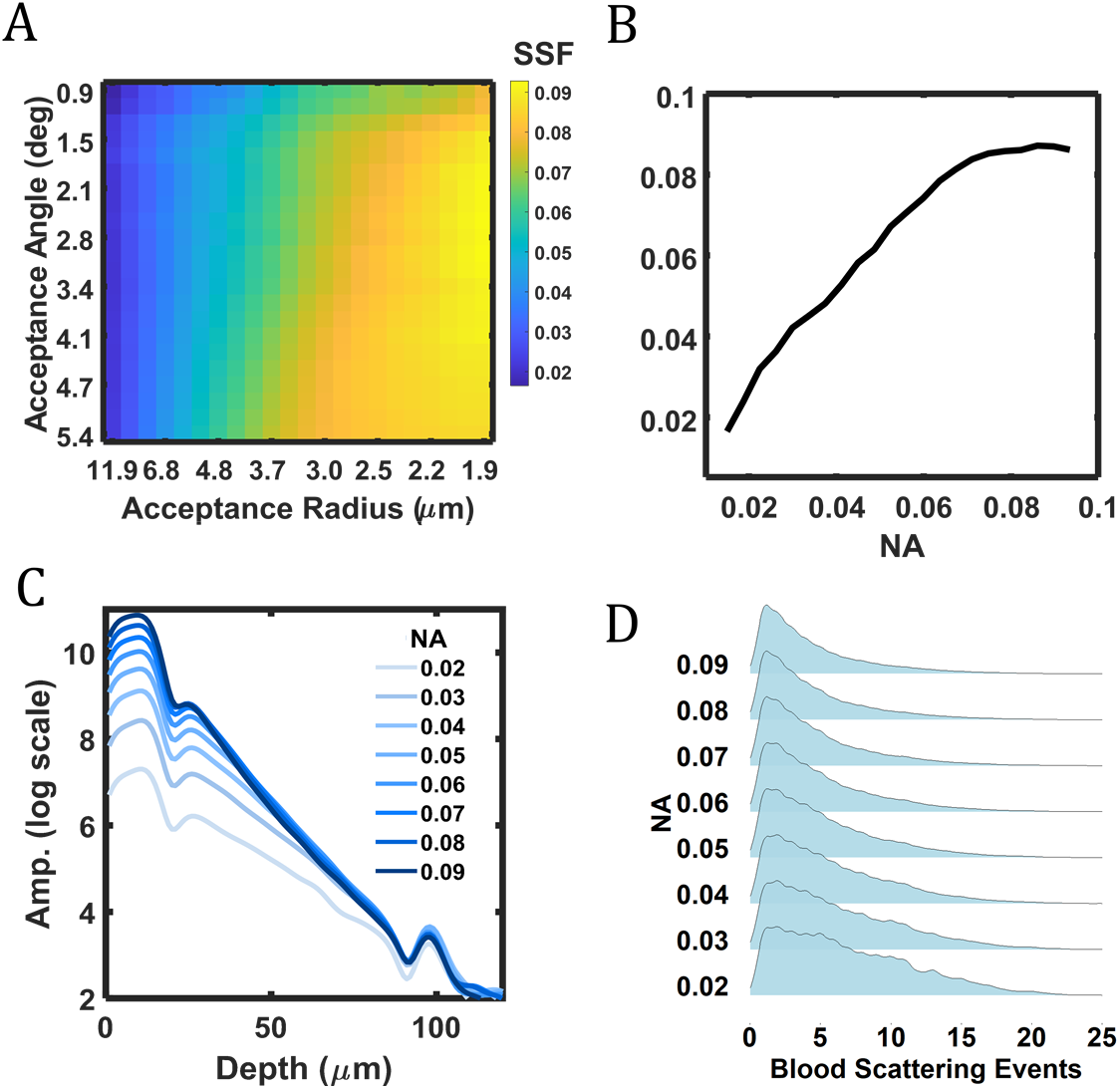
(A) The SSF values as a function of optical detection radius and angle; (B) SSF as a function of numerical aperture (NA), which is the diagonal of the matrix in the panel A; (C) Simulated A-line at center of vessel for different NAs; lighter shades indicate lower NA; (D) Normalized distributions of scattering events in blood for different

Fig. 6B plots the SSF values for physically possible NA values, which are along the diagonal of the matrix plotted in Fig. 6A, and shows that SSF increases with increased NA. However, the relationship between NA and SSF is nonlinear and appears to saturate at the highest NA. For all the tested NA values, SSF has always been less than 0.1, which is still less than one third of the literature value *W* = 0.3025. Fig. 6C plots the simulated A-line under different NA values. As NA increases, the amplitudes corresponding to AW and BM increase with respect to PW, which intrinsically increases the slope of the blood decay and leads to an increased SSF. Since the number of scattering events increases with imaging depth (Fig. 3E), photon packets propagating deeper are less likely to be detected regardless of the NA values. On the other hand, photon packets with shorter propagating depths experienced fewer scattering events and are more likely to be detected under all NA values. This explains why NA influences A-line amplitude more at shorter depths than deeper depths, as shown in Fig. 6C.

Fig. 6D plots the fitted histograms of scattering events in blood from detected photon packets for different NA values, where each histogram is normalized by its respective maximum value. Fig. 6D confirms our previous hypothesis of increased detection of multiply scattered photon packets in Figs. 6A–6C, where the histogram broadens with reduced NA values. Since the optical properties of blood remain unchanged, this difference is contributed by the changing detection criteria associated with NA.

It is important to note that this simulation presents a simplified view of the influence of OCT detection on SSF. Other variables, including the Gaussian beam profile, longitudinal chromatic aberrations, lateral chromatic aberrations, defocusing, eye geometry, oblique incidence, etc., will collectively influence the illumination and detection criteria as well [65]. However, the above simulation on the relationship between acceptance aperture and angle and SSF establishes a critical foundation for multiple scattering analysis in OCT. Such a relationship suggests no ‘one-size-fits-all’ SSF value exists.

### *Ex vivo* experimental results

Fig. 7 plots experimental vis-OCT measurements of *ex vivo* blood phantoms. All A-line reconstruction included correction for roll-off and background biases as previously reported [64, 66]. Fig. 7A shows a representative B-scan image of the phantom with fully oxygenated (sO_2_ = 100%) blood at a hematocrit of 45%, the same as our simulation. The yellow dashed line highlights the location of the A-line plotted in Fig. 7B. The A-line is an average of STFT A-lines from 528 nm – 588 nm (central band centered at 558 nm). In Fig. 7B, there is a clear delineation between the AW (40 *μ*m depth) and BM (57 *μ*m depth), corresponding to the CFZ. Attenuation measurement of this A-line using *z*_0_ = 17 *μ*m and Δ*z* = 33 *μ*m, the same as our reference depths in the simulation, gives *μ*_*sHbO*_2__(*avg*) = 226 cm^-1^ and SSF = 0.066, agreeing with the simulation.

**Fig. 7.**
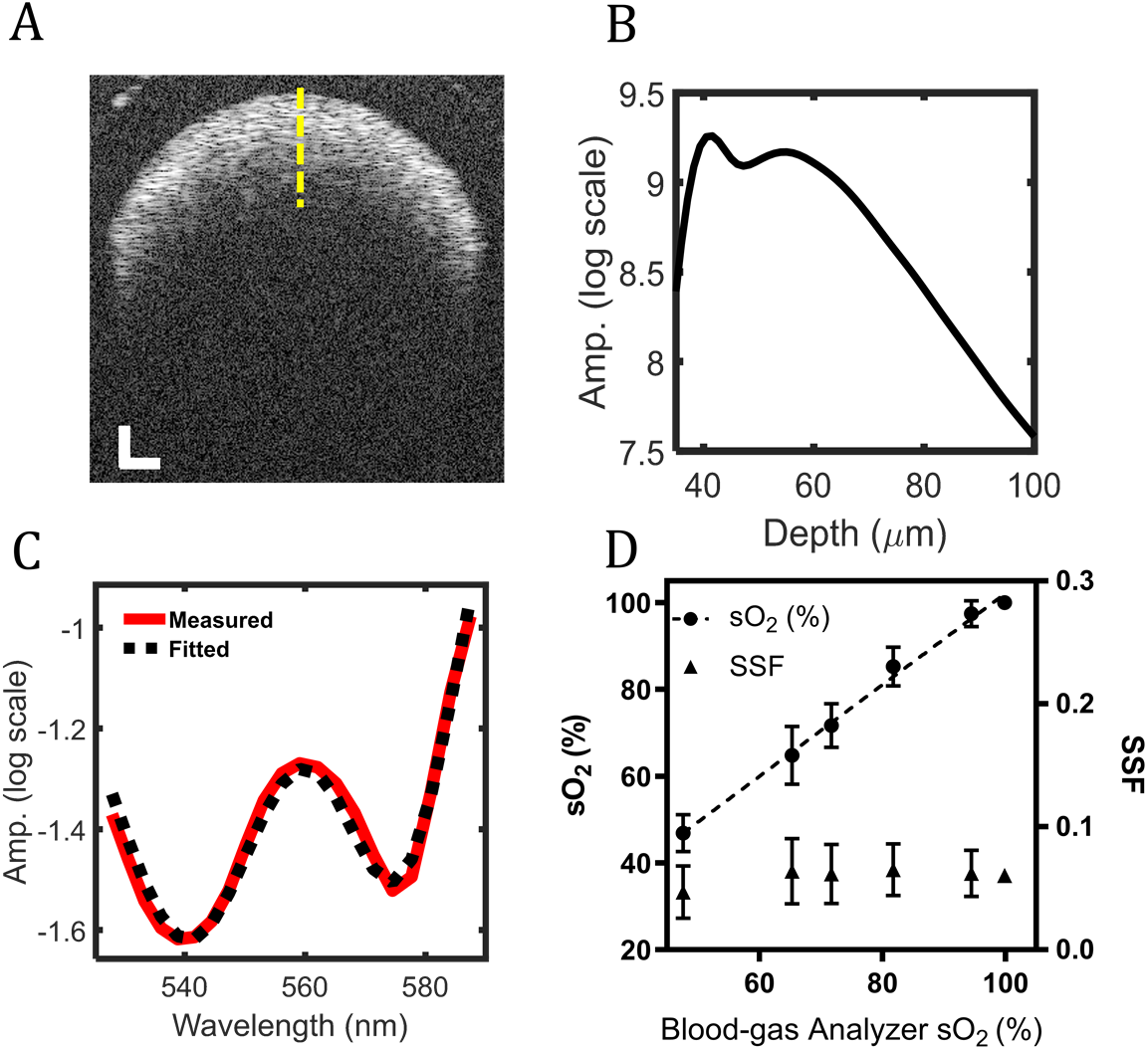
(A) vis-OCT B-scan of fully oxygenated blood in a tube phantom; (B) A vis-OCT A-line at the center of tube phantom (yellow-dashed line) in the panel A; (C) Measured attenuation spectrum of fully oxygenated blood; (D) Measured sO_2_ and SSF values at different pre-set oxygenation levels.

Fig. 7C plots a least-squares fit of the measured attenuation spectrum of fully oxygenated blood (red line) to its theoretical attenuation spectrum (black-dashed line). We performed leastsquares fitting using Eq. 4 to measure sO_2_ using the ratio of the oxygen-dependent coefficients [64] and estimate the SSF. The fitting (R^2^ = 0.98) in Fig. 7C yields sO_2_ = 100%, *μ*_*sHbO*_2__(*avg*) = 205 cm^-1^, and SSF = 0.06, which agrees with the simulated SSF for fully oxygenated blood.

We measured sO_2_ and SSF from five sO_2_ levels between 40% and 100%. We bootstrapped 100 measurements for each level by shuffling each dataset and randomly selecting 50 B-scans. Fig. 7D shows the mean and SD of measured sO_2_ (circles) and SSF (triangles). The black dashed line shows the linear best fit relationship (*y* = 1.05*x* − 3.30) between the blood-gas machine measurements and vis-OCT measurements, indicating excellent agreement between these two independent measurements. The average SSF is 0.060 +/- 0.021, and the average spectroscopic fit R^2^ is 0.99. The agreement between simulated and experimentally measured SSF values suggests that our work does not contradict the previously suggested packing factor in whole blood [9] but, instead, adds an additional correction for vis-OCT oximetry.

### *In vivo* experimental results

We validated SSF measurement in vis-OCT imaging of human retinas. Fig. 8A shows a B-scan image from a 23-year-old male volunteer. We investigated one major vein and one major artery as highlighted by 1 and 2, respectively. The yellow dashed lines highlight the locations of Alines plotted in Figs. 8B & 8C. The A-lines are an average of STFT A-lines from 528 nm – 588 nm. The vein plotted in Fig. 8B clearly delineated the AW and BM, similar to the simulated Aline (Fig. 4A) and ex vivo A-line (Fig. 7B). In the artery plotted in Fig. 8C, this delineation is less obvious, and there is a change in slope near 260 *μ*m depth where the BM is typically located, which can be caused by higher, pulsatile blood flow in arteries leading to less precise spatial averaging. We also observe a small valley near the center of the vessel in Fig. 8C, which may also be associated with more turbulent flow patterns in arteries. Nevertheless, the measured optical properties of blood also agreed with our simulated and e*x vivo* experimental results. Fig. 8D shows a least-squares fit of the attenuation spectrum measured in the vein. The best fit yields sO_2_ = 59% (R^2^ = 0.99) and SSF = 0.07. Fig. 8E shows the fitting results for the artery, where sO_2_ = 100% (R^2^ = 0.97) and SSF = 0.07.

**Fig. 8.**
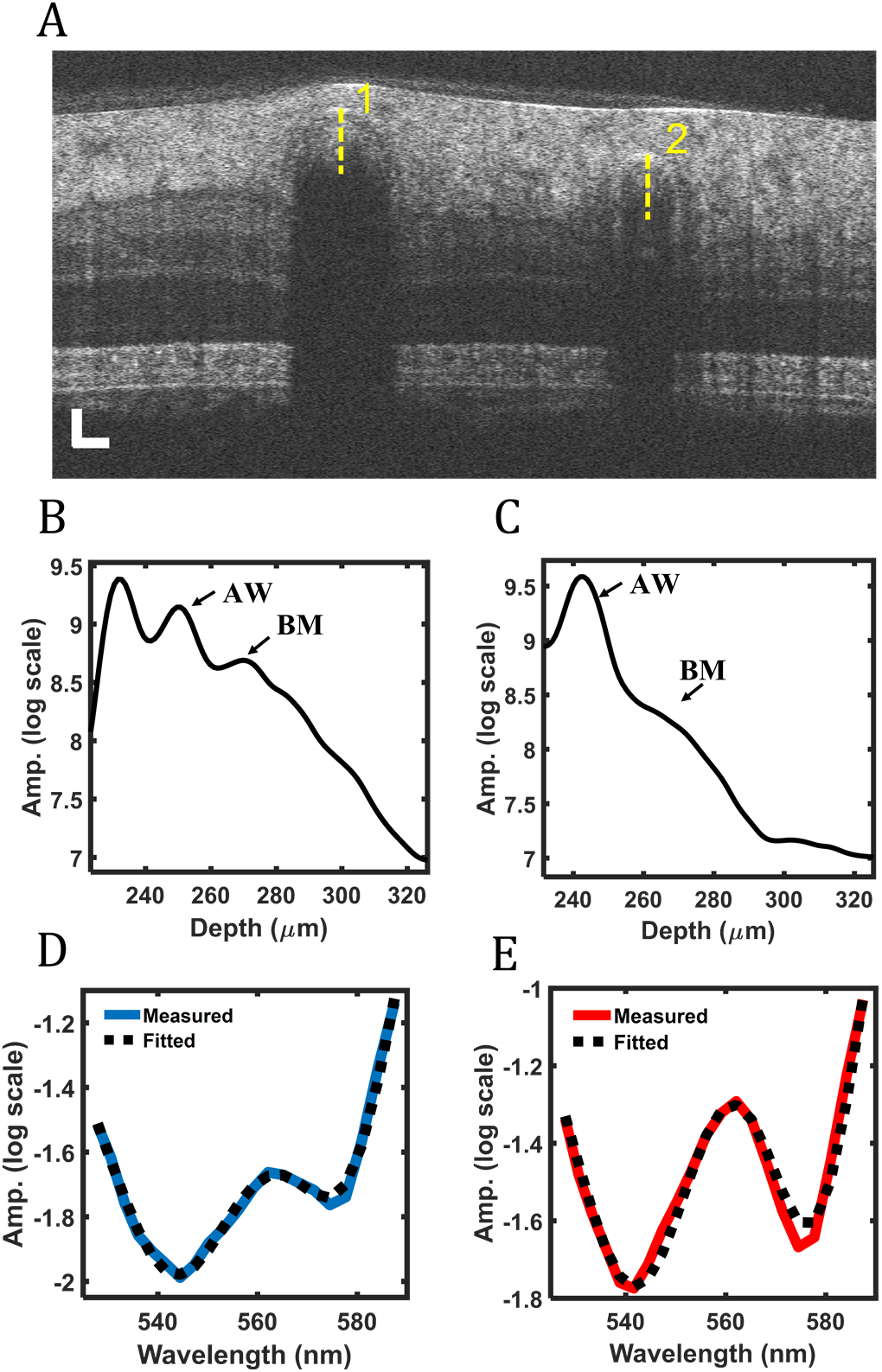
(A) Vis-OCT B-scan image of a human retina, where a vein is labeled by 1 and an artery is labeled by 2; (B) Vis-OCT A-line of the vein from the highlighted position in the panel A; (C) Vis-OCT A-line of the artery from the highlighted position in the panel (A); (D) Measured attenuation spectrum from the highlighted vein in the panel A; (E) Measured attenuation spectrum from the highlighted artery in the panel A.

## Discussion and conclusion

Accurate vis-OCT oximetry remains challenging because of systemic uncertainties introduced by multiple scattering. This work establishes a new correction factor SSF to account for multiply forward scattered photons in blood. Using MC simulation, we found that the SSF is near 0.06, significantly smaller than the reported packing factor (*W* = 0.3025) [9, 31, 55]. Documented photon packets’ trajectories indicate that most detected photon packets belong to Class I, meaning they are ballistic photon packets and return to the detector with minimal deviation from their incident axis. We found that these Class I photon packets were scattered multiple times, which is only possible if they are primarily forward scattered. We further investigated how different NAs influence the measured SSF. We found that physically reasonable NA values in the human retinal imaging yield SSF values between 0.02 and 0.09, suggesting that NA substantially impacts the measured SSF. As a result, vis-OCT should not use rigid, *a priori* models for retinal oximetry when researchers use different system hardware designs and image different eyes.

We imaged *ex vivo* bovine blood phantoms using vis-OCT as the first validation of our simulation findings. After correcting systemic biases from the background and roll-off, we measured sO_2_ and SSF and compared vis-OCT measurements with blood-oxygen analyzer measurements. We found that the average SSF was 0.060 +/- 0.021, almost identical to the simulated results. We further performed vis-OCT sO_2_ measurements in the human retina and found that SSF was 0.06 and measured sO_2_ values consistent with physiological ranges for arteries and veins. This work is the first comprehensive investigation and validation of blood’s attenuation spectrum in vis-OCT using simulation, *ex vivo* phantom imaging, and human retinal imaging.

Using the packing factor to scale the scattering coefficient in our MC simulation resulted in SSF values in excellent agreement with experimental data. As suggested in the literature, the packing factor is the result of correlated scatterings among densely packed RBCs [9, 29, 67, 68]. As RBC concentration increases, coherent interferences can affect the far-field scattered field, which is nonlinearly correlated with RBC density and is likely dependent on the orientations of individual RBCs. While MC simulation does not directly account for orientation-dependent scattering, it can replicate their statistical influence by scaling the input *μ*_s_ by *W*. Furthermore, previous experimental tests of blood’s scattering coefficient were performed using an integration sphere [10, 11, 29, 47, 69], which does not spatially filter detected photons as vis-OCT does. Our SSF combines the influence of scattering effects from blood hemodynamics (e.g., packing factor) and spatial filtering by the imaging modality (e.g., acceptance aperture and angle) on the effective *μ*_s_ measured by vis-OCT.

One limitation in our simulation is the assumption of blood as a homogenous medium. In reality, RBC packing density and orientations are affected by blood flow, blood velocity, vessel size, and incident angle [11, 50, 70–72]. These factors can alter scattering cross-section and directionality, changing the optical properties assumed in this work. Although significant spatial averaging may suppress these variations, researchers should carefully monitor the variability of SSF to ensure a suitable oximetry model.

We investigated a suite of parameters influencing vis-OCT oximetry. We found that the combination of a low NA and high forward scattering breaks the common belief that OCT is mainly sensitive to blood’s scattering coefficient because ophthalmic vis-OCT almost exclusively detects the unique Class I photons in the blood. Furthermore, considering the contribution of SSF, the detected scattering coefficient is significantly lower than the expected scattering coefficient, allowing absorption-dominated measurements in vis-OCT oximetry.

## Acknowledgment

We acknowledge the generous financial support from NIH grants R01EY026078, R01EY019949, R01EY029121, U01EY033001, and T32GM142604.

## Notes

### Competing Interest Statement

The authors have declared no competing interest.

